## Supplemental Material for "Heterotypic stressors unmask behavioral influences of PMAT deficiency in mice"

Supplemental Table S1. Repeated measures ANOVAs of Phase 1 Cued Female data.

### Phase 1 Cued Females

#### Training

| | *F statistic* | *p* | *partial $\eta^2$* |
|---|---|---|---|
| Genotype | F(1,35)=0.531 | 0.471 | 0.015 |
| Swim | F(1,35)=0.003 | 0.957 | 0.000 |
| Genotype × Swim | F(1,35)=0.881 | 0.354 | 0.025 |
| **Time** | **F(3.122,109.266)=183.4** | **<0.001** | **0.840** |
| Time × Genotype | F(3.122,109.266)=0.676 | 0.574 | 0.019 |
| Time × Swim | F(3.122,109.266)=0.585 | 0.633 | 0.016 |
| Time × Genotype × Swim | F(3.122,109.266)=0.964 | 0.415 | 0.027 |

#### Expression Testing & Extinction Training

| | *F statistic* | *p* | *partial $\eta^2$* |
|---|---|---|---|
| Genotype | F(1,35)=0.471 | 0.497 | 0.013 |
| Swim | F(1,35)=0.455 | 0.504 | 0.013 |
| Genotype × Swim | F(1,35)=0.751 | 0.392 | 0.021 |
| Time | F(9.728,340.487)=8.714 | <0.001 | 0.199 |
| **Time × Genotype** | **F(9.728,340.487)=3.865** | **<0.001** | **0.099** |
| Time × Swim | F(9.728,340.487)=0.802 | 0.624 | 0.022 |
| Time × Genotype × Swim | F(9.728,340.487)=1.138 | 0.334 | 0.031 |

#### Extinction Retention Testing & More Extinction Training

| | *F statistic* | *p* | *partial $\eta^2$* |
|---|---|---|---|
| Genotype | F(1,35)=1.000 | 0.324 | 0.028 |
| Swim | F(1,35)=0.477 | 0.494 | 0.013 |
| Genotype × Swim | F(1,35)=0.508 | 0.481 | 0.014 |
| **Time** | **F(9.774,342.097)=8.024** | **<0.001** | **0.186** |
| Time × Genotype | F(9.774,342.097)=1.034 | 0.414 | 0.029 |
| Time × Swim | F(9.774,342.097)=1.523 | 0.131 | 0.042 |
| Time × Genotype × Swim | F(9.774,342.097)=1.191 | 0.296 | 0.033 |

#### Context Fear Expression - Timecourse

| | *F statistic* | *p* | *partial $\eta^2$* |
|---|---|---|---|
| Genotype | F(1,35)=0.001 | 0.976 | 0.000 |
| Swim | F(1,35)=0.110 | 0.742 | 0.003 |
| Genotype × Swim | F(1,35)=0.027 | 0.871 | 0.001 |
| **Time** | **F(6.926,242.403)=5.297** | **<0.001** | **0.131** |
| Time × Genotype | F(6.926,242.403)=0.641 | 0.720 | 0.018 |
| Time × Swim | F(6.926,242.403)=0.586 | 0.765 | 0.016 |

| | | | |
|---|---|---|---|
| Time × Genotype × Swim | F(6.926,242.403)=0.531 | 0.809 | 0.015 |

Cued Fear Renewal

| | ***F statistic*** | ***p*** | ***partial η²*** |
|---|---|---|---|
| Genotype | F(1,35)=0.009 | 0.926 | 0.000 |
| Swim | F(1,35)=0.075 | 0.786 | 0.002 |
| Genotype × Swim | F(1,35)=0.421 | 0.521 | 0.012 |
| **Time** | **F(3.712,129.923)=12.40** | **<0.001** | **0.262** |
| Time × Genotype | F(3.712,129.923)=0.574 | 0.670 | 0.016 |
| Time × Swim | F(3.712,129.923)=0.745 | 0.554 | 0.021 |
| Time × Genotype × Swim | F(3.712,129.923)=0.184 | 0.937 | 0.005 |

Supplemental Table 2. Repeated measures ANOVAs of Phase 1 Cued Male data.

### Phase 1 Cued Males

#### Training

| | *F statistic* | *p* | *partial $\eta^2$* |
|---|---|---|---|
| Genotype | F(1,54)=3.148 | 0.082 | 0.055 |
| Swim | F(1,54)=1.825 | 0.182 | 0.033 |
| Genotype × Swim | F(1,54)=0.166 | 0.685 | 0.003 |
| **Time** | **F(3.156,170.433)=205.7** | **<0.001** | **0.792** |
| Time × Genotype | F(3.156,170.433)=1.554 | 0.200 | 0.028 |
| Time × Swim | F(3.156,170.433)=0.475 | 0.710 | 0.009 |
| Time × Genotype × Swim | F(3.156,170.433)=0.477 | 0.708 | 0.009 |

#### Expression Testing & Extinction Training

| | *F statistic* | *p* | *partial $\eta^2$* |
|---|---|---|---|
| Genotype | F(1,54)=1.051 | 0.310 | 0.019 |
| Swim | F(1,54)=0.074 | 0.787 | 0.001 |
| Genotype × Swim | F(1,54)=2.257 | 0.139 | 0.040 |
| **Time** | **F(10.594,572.057)=9.786** | **<0.001** | **0.153** |
| Time × Genotype | F(10.594,572.057)=0.538 | 0.872 | 0.010 |
| Time × Swim | F(10.594,572.057)=0.554 | 0.860 | 0.010 |
| Time × Genotype × Swim | F(10.594,572.057)=0.718 | 0.716 | 0.013 |

#### Extinction Retention Testing & More Extinction Training

| | *F statistic* | *p* | *partial $\eta^2$* |
|---|---|---|---|
| Genotype | F(1,53)=0.173 | 0.679 | 0.003 |
| Swim | F(1,53)=0.035 | 0.852 | 0.001 |
| Genotype × Swim | F(1,53)=1.543 | 0.220 | 0.028 |
| **Time** | **F(10.125,536.64)=11.09** | **<0.001** | **0.173** |
| Time × Genotype | F(10.125,536.64)=1.004 | 0.439 | 0.019 |
| Time × Swim | F(10.125,536.64)=1.339 | 0.205 | 0.025 |
| Time × Genotype × Swim | F(10.125,536.64)=0.633 | 0.788 | 0.012 |

#### Context Fear Expression

| | *F statistic* | *p* | *partial $\eta^2$* |
|---|---|---|---|
| Genotype | F(1,49)=0.250 | 0.619 | 0.005 |
| Swim | F(1,49)=0.684 | 0.412 | 0.014 |
| Genotype × Swim | F(1,49)=1.147 | 0.289 | 0.023 |
| **Time** | **F(10.889,533.558)=11.58** | **<0.001** | **0.191** |
| Time × Genotype | F(10.889,533.558)=0.631 | 0.801 | 0.013 |
| Time × Swim | F(10.889,533.558)=1.197 | 0.286 | 0.024 |

| | | | |
|---|---|---|---|
| Time × Genotype × Swim | F(10.889,533.558)=1.319 | 0.210 | 0.026 |

Cued Fear Renewal

| | ***F statistic*** | ***p*** | ***partial η²*** |
|---|---|---|---|
| Genotype | F(1,49)=0.033 | 0.856 | 0.001 |
| Swim | F(1,49)=0.874 | 0.354 | 0.018 |
| Genotype × Swim | F(1,49)=0.081 | 0.777 | 0.002 |
| **Time** | **F(3.82,187.198)=11.17** | **<0.001** | **0.186** |
| Time × Genotype | F(3.82,187.198)=0.190 | 0.938 | 0.004 |
| Time × Swim | F(3.82,187.198)=0.182 | 0.942 | 0.004 |
| Time × Genotype × Swim | F(3.82,187.198)=0.999 | 0.407 | 0.020 |

Supplemental Table 3. Repeated measures ANOVAs of Phase 1 Context Female data.

### Phase 1 Context Females

#### Training

| | *F statistic* | *p* | *partial $\eta^2$* |
|---|---|---|---|
| Genotype | F(1,34)=0.904 | 0.348 | 0.026 |
| Swim | F(1,34)=0.014 | 0.907 | 0.000 |
| Genotype × Swim | F(1,34)=0.061 | 0.806 | 0.002 |
| **Time** | **F(3.626,123.285)=125.3** | **<0.001** | **0.787** |
| Time × Genotype | F(3.626,123.285)=0.252 | 0.893 | 0.007 |
| Time × Swim | F(3.626,123.285)=0.818 | 0.506 | 0.023 |
| Time × Genotype × Swim | F(3.626,123.285)=0.412 | 0.782 | 0.012 |

#### Context Fear Expression

| | *F statistic* | *p* | *partial $\eta^2$* |
|---|---|---|---|
| *Genotype* | *F(1,34)=2.936* | *0.096* | *0.079* |
| Swim | F(1,34)=0.540 | 0.468 | 0.016 |
| Genotype × Swim | F(1,34)=0.555 | 0.462 | 0.016 |
| **Time** | **F(9.138,310.686)=8.874** | **<0.001** | **0.207** |
| Time × Genotype | F(9.138,310.686)=1.369 | 0.200 | 0.039 |
| Time × Swim | F(9.138,310.686)=0.494 | 0.881 | 0.014 |
| Time × Genotype × Swim | F(9.138,310.686)=0.984 | 0.454 | 0.028 |

Supplemental Table 4.  Repeated measures ANOVAs of Phase 1 Context Male data.

| **Phase 1 Context Males** | | | |
|---|---|---|---|
| Training | | | |
| | ***F statistic*** | ***p*** | ***partial η²*** |
| Genotype | F(1,39)=0.553 | 0.462 | 0.014 |
| Swim | F(1,39)=2.221 | 0.144 | 0.054 |
| Genotype × Swim | F(1,39)=0.081 | 0.777 | 0.002 |
| **Time** | **F(3.226,125.803)=12.92** | **<0.001** | **0.249** |
| Time × Genotype | F(3.226,125.803)=1.424 | 0.237 | 0.035 |
| Time × Swim | F(3.226,125.803)=0.441 | 0.738 | 0.011 |
| Time × Genotype × Swim | F(3.226,125.803)=0.158 | 0.934 | 0.004 |
| Context Fear Expression | | | |
| | ***F statistic*** | ***p*** | ***partial η²*** |
| **Genotype** | **F(1,39)=4.555** | **0.039** | **0.105** |
| Swim | F(1,39)=0.027 | 0.871 | 0.001 |
| Genotype × Swim | F(1,39)=1.280 | 0.265 | 0.032 |
| **Time** | **F(9.546,372.282)=13.94** | **<0.001** | **0.263** |
| Time × Genotype | F(9.546,372.282)=1.142 | 0.331 | 0.028 |
| Time × Swim | F(9.546,372.282)=1.319 | 0.221 | 0.033 |
| Time × Genotype × Swim | F(9.546,372.282)=0.840 | 0.586 | 0.021 |

Supplemental Table 5. Repeated measures ANOVAs of Phase 2 Cued Female data.

### Phase 2 Cued Females

#### Training

| | ***F statistic*** | ***p*** | ***partial η²*** |
|---|---|---|---|
| Genotype | F(1,41)=0.676 | 0.416 | 0.016 |
| Swim | F(1,41)=1.273 | 0.266 | 0.030 |
| Genotype × Swim | F(1,41)=0.922 | 0.343 | 0.022 |
| **Time** | **F(3.605,147.811)=159.1** | **<0.001** | **0.795** |
| Time × Genotype | F(3.605,147.811)=0.679 | 0.593 | 0.016 |
| Time × Swim | F(3.605,147.811)=0.970 | 0.420 | 0.023 |
| Time × Genotype × Swim | F(3.605,147.811)=0.169 | 0.942 | 0.004 |

#### Expression Testing & Extinction Training

| | ***F statistic*** | ***p*** | ***partial η²*** |
|---|---|---|---|
| Genotype | F(1,41)=0.299 | 0.587 | 0.007 |
| Swim | F(1,41)=0.036 | 0.850 | 0.001 |
| Genotype × Swim | F(1,41)=0.951 | 0.335 | 0.023 |
| **Time** | **F(8.741,358.393)=6.623** | **<0.001** | **0.139** |
| Time × Genotype | F(8.741,358.393)=0.852 | 0.566 | 0.020 |
| Time × Swim | F(8.741,358.393)=0.726 | 0.681 | 0.017 |
| Time × Genotype × Swim | F(8.741,358.393)=0.437 | 0.911 | 0.011 |

#### Extinction Retention Testing & More Extinction Training

| | ***F statistic*** | ***p*** | ***partial η²*** |
|---|---|---|---|
| Genotype | F(1,35)=0.400 | 0.531 | 0.011 |
| Swim | F(1,35)=0.093 | 0.763 | 0.003 |
| Genotype × Swim | F(1,35)=0.009 | 0.924 | 0.000 |
| **Time** | **F(8.864,310.252)=11.15** | **<0.001** | **0.242** |
| Time × Genotype | F(8.864,310.252)=0.590 | 0.803 | 0.017 |
| Time × Swim | F(8.864,310.252)=0.822 | 0.594 | 0.023 |
| Time × Genotype × Swim | F(8.864,310.252)=1.133 | 0.339 | 0.031 |

#### Context Fear Expression

| | ***F statistic*** | ***p*** | ***partial η²*** |
|---|---|---|---|
| Genotype | F(1,41)=0.256 | 0.615 | 0.006 |
| Swim | F(1,41)=0.498 | 0.484 | 0.012 |
| Genotype × Swim | F(1,41)=0.060 | 0.807 | 0.001 |
| **Time** | **F(10.762,441.257)=5.923** | **<0.001** | **0.126** |
| Time × Genotype | F(10.762,441.257)=1.001 | 0.444 | 0.024 |
| Time × Swim | F(10.762,441.257)=1.15 | 0.321 | 0.027 |

| | | | |
|---|---|---|---|
| Time × Genotype × Swim | F(10.762,441.257)=1.064 | 0.389 | 0.025 |

Cued Fear Renewal

| | *F statistic* | *p* | *partial* $\eta^2$ |
|---|---|---|---|
| Genotype | F(1,39)=0.637 | 0.430 | 0.016 |
| Swim | F(1,39)=0.050 | 0.825 | 0.001 |
| Genotype × Swim | F(1,39)=0.066 | 0.799 | 0.002 |
| Time | F(3.57,139.219)=11.74 | <0.001 | 0.231 |
| Time × Genotype | F(3.57,139.219)=0.690 | 0.584 | 0.017 |
| **Time × Swim** | **F(3.57,139.219)=3.592** | **0.011** | **0.084** |
| Time × Genotype × Swim | F(3.57,139.219)=0.282 | 0.870 | 0.007 |

Supplemental Table 6. Repeated measures ANOVAs of Phase 2 Cued Male data.

### Phase 2 Cued Males

#### Training

| | *F statistic* | *p* | *partial* $\eta^2$ |
|---|---|---|---|
| Genotype | F(1,36)=0.033 | 0.858 | 0.001 |
| Swim | F(1,36)=0.140 | 0.710 | 0.004 |
| Genotype × Swim | F(1,36)=1.226 | 0.276 | 0.033 |
| **Time** | **F(2.700,97.197)=123.1** | **<0.001** | **0.774** |
| Time × Genotype | F(2.700,97.197)=1.581 | 0.203 | 0.042 |
| Time × Swim | F(2.700,97.197)=0.872 | 0.449 | 0.024 |
| Time × Genotype × Swim | F(2.700,97.197)=0.978 | 0.400 | 0.026 |

#### Expression Testing & Extinction Training

| | *F statistic* | *p* | *partial* $\eta^2$ |
|---|---|---|---|
| Genotype | F(1,30)=0.196 | 0.661 | 0.006 |
| Swim | F(1,30)=1.649 | 0.209 | 0.052 |
| Genotype × Swim | F(1,30)=0.725 | 0.401 | 0.024 |
| **Time** | **F(8.933,267.996)=10.55** | **<0.001** | **0.260** |
| Time × Genotype | F(8.933,267.996)=1.029 | 0.417 | 0.033 |
| Time × Swim | F(8.933,267.996)=0.987 | 0.451 | 0.032 |
| Time × Genotype × Swim | F(8.933,267.996)=1.631 | 0.107 | 0.052 |

#### Extinction Retention Testing & More Extinction Training

| | *F statistic* | *p* | *partial* $\eta^2$ |
|---|---|---|---|
| **Genotype** | **F(1,38)=6.914** | **0.012** | **0.154** |
| Swim | F(1,38)=0.052 | 0.820 | 0.001 |
| Genotype × Swim | F(1,38)=1.033 | 0.316 | 0.026 |
| **Time** | **F(7.79,296.005)=8.583** | **<0.001** | **0.184** |
| Time × Genotype | F(7.79,296.005)=0.955 | 0.470 | 0.025 |
| Time × Swim | F(7.79,296.005)=1.211 | 0.293 | 0.031 |
| Time × Genotype × Swim | F(7.79,296.005)=1.249 | 0.272 | 0.032 |

#### Context Fear Expression

| | *F statistic* | *p* | *partial* $\eta^2$ |
|---|---|---|---|
| **Genotype** | **F(1,37)=4.175** | **0.048** | **0.101** |
| Swim | F(1,37)=0.273 | 0.604 | 0.007 |
| Genotype × Swim | F(1,37)=2.728 | 0.107 | 0.069 |
| **Time** | **F(9.633,356.405)=5.218** | **<0.001** | **0.124** |
| Time × Genotype | F(9.633,356.405)=0.390 | 0.947 | 0.010 |
| Time × Swim | F(9.633,356.405)=1.470 | 0.152 | 0.038 |

| | | | |
|---|---|---|---|
| Time × Genotype × Swim | F(9.633,356.405)=1.183 | 0.302 | 0.031 |

Cued Fear Renewal

| | *F statistic* | *p* | *partial η²* |
|---|---|---|---|
| Genotype | F(1,34)=0.164 | 0.688 | 0.005 |
| Swim | F(1,34)=1.950 | 0.172 | 0.054 |
| Genotype × Swim | F(1,34)=0.625 | 0.435 | 0.018 |
| **Time** | **F(3.835,130.383)=24.30** | **<0.001** | **0.417** |
| Time × Genotype | F(3.835,130.383)=0.851 | 0.492 | 0.024 |
| Time × Swim | F(3.835,130.383)=0.898 | 0.464 | 0.026 |
| Time × Genotype × Swim | F(3.835,130.383)=1.335 | 0.262 | 0.038 |

Supplemental Table 7. Repeated measures ANOVAs of Phase 2 Context Female data.

**Phase 2 Context Females**

Training

| | ***F statistic*** | ***p*** | ***partial η²*** |
|---|---|---|---|
| Genotype | F(1,34)=0.713 | 0.404 | 0.021 |
| Swim | F(1,34)=0.346 | 0.560 | 0.010 |
| Genotype × Swim | F(1,34)=0.065 | 0.800 | 0.002 |
| **Time** | **F(3.029,102.984)=76.52** | **<0.001** | **0.692** |
| Time × Genotype | F(3.029,102.984)=1.116 | 0.346 | 0.032 |
| Time × Swim | F(3.029,102.984)=1.963 | 0.124 | 0.055 |
| Time × Genotype × Swim | F(3.029,102.984)=0.978 | 0.407 | 0.028 |

Context Fear Expression

| | ***F statistic*** | ***p*** | ***partial η²*** |
|---|---|---|---|
| Genotype | F(1,30)=0.362 | 0.552 | 0.012 |
| Swim | F(1,30)=2.066 | 0.161 | 0.064 |
| Genotype × Swim | F(1,30)=1.549 | 0.223 | 0.049 |
| **Time** | **F(9.209,276.265)=8.773** | **<0.001** | **0.226** |
| Time × Genotype | F(9.209,276.265)=1.094 | 0.367 | 0.035 |
| Time × Swim | F(9.209,276.265)=0.985 | 0.454 | 0.032 |
| Time × Genotype × Swim | F(9.209,276.265)=1.251 | 0.263 | 0.040 |

Supplemental Table 8. Repeated measures ANOVAs of Phase 2 Context Male data.

**Phase 2 Context Males**

Training

| | *F statistic* | *p* | *partial η²* |
|---|---|---|---|
| Genotype | F(1,33)=0.042 | 0.839 | 0.001 |
| Swim | F(1,33)=0.075 | 0.786 | 0.002 |
| Genotype × Swim | F(1,33)=0.180 | 0.674 | 0.005 |
| **Time** | **F(3.284,108.38)=66.40** | **<0.001** | **0.668** |
| Time × Genotype | F(3.284,108.38)=0.190 | 0.917 | 0.006 |
| Time × Swim | F(3.284,108.38)=0.979 | 0.411 | 0.029 |
| Time × Genotype × Swim | F(3.284,108.38)=1.062 | 0.372 | 0.031 |

Context Fear Expression

| | *F statistic* | *p* | *partial η²* |
|---|---|---|---|
| Genotype | F(1,33)=0.524 | 0.474 | 0.016 |
| Swim | F(1,33)=0.024 | 0.877 | 0.001 |
| *Genotype × Swim* | *F(1,33)=3.400* | *0.074* | *0.093* |
| **Time** | **F(8.625,284.634)=14.87** | **<0.001** | **0.311** |
| Time × Genotype | F(8.625,284.634)=1.447 | 0.171 | 0.042 |
| Time × Swim | F(8.625,284.634)=1.160 | 0.322 | 0.034 |
| Time × Genotype × Swim | F(8.625,284.634)=0.763 | 0.646 | 0.023 |

Supplemental Table 9. Three-way ANOVAs of log-transformed corticosterone levels taken 2 h following final testing (Phase 1 – swim stress; Phase 2 – context testing & cued fear renewal).

| **Phase 1 Cued** | | | |
|---|---|---|---|
| | ***F statistic*** | ***p*** | ***partial $\eta^2$*** |
| Genotype | F(1,82)=0.962 | 0.330 | 0.012 |
| **Sex** | **F(1,82)=17.77** | **<0.001** | **0.178** |
| Swim | F(1,82)=0.301 | 0.584 | 0.004 |
| Genotype × Sex | F(1,82)=0.119 | 0.731 | 0.001 |
| Genotype × Swim | F(1,82)=0.170 | 0.681 | 0.002 |
| Sex × Swim | F(1,82)=0.000 | 0.992 | 0.000 |
| Genotype × Sex × Swim | F(1,82)=0.639 | 0.426 | 0.008 |

| **Phase 1 Context** | | | |
|---|---|---|---|
| | ***F statistic*** | ***p*** | ***partial $\eta^2$*** |
| Genotype | F(1,72)=2.007 | 0.161 | 0.027 |
| Sex | F(1,72)=45.11 | <0.001 | 0.385 |
| Swim | F(1,72)=8.382 | 0.005 | 0.104 |
| Genotype × Sex | F(1,72)=4.839 | 0.031 | 0.063 |
| Genotype × Swim | F(1,72)=0.239 | 0.626 | 0.003 |
| Sex × Swim | F(1,72)=5.938 | 0.017 | 0.076 |
| **Genotype × Sex × Swim** | **F(1,72)=6.029** | **0.016** | **0.077** |

| **Phase 2 Cued** | | | |
|---|---|---|---|
| | ***F statistic*** | ***p*** | ***partial $\eta^2$*** |
| Genotype | F(1,79)=2.285 | 0.135 | 0.028 |
| **Sex** | **F(1,79)=17.40** | **<0.001** | **0.180** |
| Swim | F(1,79)=0.967 | 0.328 | 0.012 |
| Genotype × Sex | F(1,79)=1.157 | 0.285 | 0.014 |
| Genotype × Swim | F(1,79)=1.748 | 0.190 | 0.022 |
| Sex × Swim | F(1,79)=0.390 | 0.534 | 0.005 |
| Genotype × Sex × Swim | F(1,79)=1.235 | 0.270 | 0.015 |

| **Phase 2 Context** | | | |
|---|---|---|---|
| | ***F statistic*** | ***p*** | ***partial $\eta^2$*** |
| Genotype | F(1,63)=0.001 | 0.979 | 0.000 |
| Sex | F(1,63)=1.437 | 0.235 | 0.022 |
| Swim | F(1,63)=0.670 | 0.416 | 0.011 |
| Genotype × Sex | F(1,63)=0.186 | 0.668 | 0.003 |
| **Genotype × Swim** | **F(1,63)=4.377** | **0.040** | **0.065** |
| Sex × Swim | F(1,63)=0.100 | 0.753 | 0.002 |

| | | | |
|---|---|---|---|
| Genotype × Sex × Swim | $F(1,63)=0.282$ | 0.597 | 0.004 |

Supplemental Table 10. Two-way ANOVAs for behaviors during swim stress in Phase 1 Cued mice.

**Phase 1 Cued**

Swimming

| | ***F statistic*** | ***p*** | ***partial η²*** |
|---|---|---|---|
| Genotype | F(1,39)=0.010 | 0.922 | 0.000 |
| Sex | F(1,39)=1.825 | 0.185 | 0.045 |
| Genotype × Sex | F(1,39)=0.419 | 0.521 | 0.011 |

Immobility

| | ***F statistic*** | ***p*** | ***partial η²*** |
|---|---|---|---|
| Genotype | F(1,39)=0.801 | 0.376 | 0.020 |
| Sex | F(1,39)=0.065 | 0.801 | 0.002 |
| Genotype × Sex | F(1,39)=0.570 | 0.455 | 0.014 |

Climbing

| | ***F statistic*** | ***p*** | ***partial η²*** |
|---|---|---|---|
| Genotype | F(1,39)=0.403 | 0.529 | 0.010 |
| Sex | F(1,39)=0.626 | 0.434 | 0.016 |
| Genotype × Sex | F(1,39)=0.134 | 0.716 | 0.003 |

Latency to 1st Immobility

| | ***F statistic*** | ***p*** | ***partial η²*** |
|---|---|---|---|
| Genotype | F(1,39)=0.356 | 0.554 | 0.009 |
| Sex | F(1,39)=1.051 | 0.312 | 0.026 |
| Genotype × Sex | F(1,39)=0.003 | 0.957 | 0.000 |

Supplemental Table 11. Two-way ANOVAs for behaviors during swim stress in Phase 1 Context mice.

**Phase 1 Context**

Swimming

| | *F statistic* | *p* | *partial η²* |
|---|---|---|---|
| Genotype | F(1,36)=0.013 | 0.911 | 0.000 |
| Sex | F(1,36)=4.250 | 0.047 | 0.106 |
| **Genotype × Sex** | **F(1,36)=5.572** | **0.024** | **0.134** |

Immobility

| | *F statistic* | *p* | *partial η²* |
|---|---|---|---|
| Genotype | F(1,36)=0.283 | 0.598 | 0.008 |
| Sex | F(1,36)=0.181 | 0.673 | 0.005 |
| Genotype × Sex | F(1,36)=1.360 | 0.251 | 0.036 |

Climbing

| | *F statistic* | *p* | *partial η²* |
|---|---|---|---|
| Genotype | F(1,36)=0.765 | 0.388 | 0.021 |
| Sex | F(1,36)=1.538 | 0.223 | 0.041 |
| Genotype × Sex | F(1,36)=0.189 | 0.667 | 0.005 |

Latency to 1$^{st}$ Immobility

| | *F statistic* | *p* | *partial η²* |
|---|---|---|---|
| Genotype | F(1,36)=0.151 | 0.700 | 0.004 |
| Sex | F(1,36)=0.259 | 0.614 | 0.007 |
| Genotype × Sex | F(1,36)=2.022 | 0.164 | 0.053 |

Supplemental Table 12. Two-way ANOVAs for behaviors during swim stress in Phase 2 Cued mice.

**Phase 2 Cued**

Swimming

| | *F statistic* | *p* | *partial η²* |
|---|---|---|---|
| Genotype | F(1,42)=1.062 | 0.309 | 0.025 |
| Sex | F(1,42)=0.048 | 0.828 | 0.001 |
| Genotype × Sex | F(1,42)=0.271 | 0.605 | 0.006 |

Immobility

| | *F statistic* | *p* | *partial η²* |
|---|---|---|---|
| *Genotype* | *F(1,42)=3.221* | *0.080* | *0.071* |
| Sex | F(1,42)=0.065 | 0.800 | 0.002 |
| Genotype × Sex | F(1,42)=1.161 | 0.287 | 0.027 |

Climbing

| | *F statistic* | *p* | *partial η²* |
|---|---|---|---|
| *Genotype* | *F(1,42)=3.024* | *0.089* | *0.067* |
| Sex | F(1,42)=0.867 | 0.357 | 0.020 |
| Genotype × Sex | F(1,42)=1.483 | 0.230 | 0.034 |

Latency to 1st Immobility

| | *F statistic* | *p* | *partial η²* |
|---|---|---|---|
| **Genotype** | **F(1,42)=4.679** | **0.036** | **0.100** |
| Sex | F(1,42)=2.722 | 0.106 | 0.061 |
| Genotype × Sex | F(1,42)=0.030 | 0.863 | 0.001 |

Supplemental Table 13. Two-way ANOVAs for behaviors during swim stress in Phase 2 Context mice.

### Phase 2 Context

#### Swimming

| | *F statistic* | *p* | *partial η²* |
|---|---|---|---|
| Genotype | F(1,34)=0.694 | 0.411 | 0.020 |
| **Sex** | **F(1,34)=4.996** | **0.032** | **0.128** |
| Genotype × Sex | F(1,34)=0.003 | 0.954 | 0.000 |

#### Immobility

| | *F statistic* | *p* | *partial η²* |
|---|---|---|---|
| Genotype | F(1,34)=1.512 | 0.227 | 0.043 |
| **Sex** | **F(1,34)=10.09** | **0.003** | **0.229** |
| Genotype × Sex | F(1,34)=0.013 | 0.909 | 0.000 |

#### Climbing

| | *F statistic* | *p* | *partial η²* |
|---|---|---|---|
| Genotype | F(1,34)=0.930 | 0.342 | 0.027 |
| **Sex** | **F(1,34)=5.611** | **0.024** | **0.142** |
| Genotype × Sex | F(1,34)=0.020 | 0.890 | 0.001 |

#### Latency to 1$^{st}$ Immobility

| | *F statistic* | *p* | *partial η²* |
|---|---|---|---|
| Genotype | F(1,34)=0.845 | 0.364 | 0.024 |
| **Sex** | **F(1,34)=18.76** | **<0.001** | **0.356** |
| Genotype × Sex | F(1,34)=0.087 | 0.770 | 0.003 |

Supplemental Table S14. Two-way ANOVA of log-transformed corticosterone levels in mice 30 min following an acute swim stressor. Refer to Supplemental Figure S1.

| **Log-transformed Corticosterone Levels** | | | |
|---|---|---|---|
| | ***F statistic*** | ***p*** | ***partial*** $\eta^2$ |
| Genotype | F(2,62)=0.285 | 0.753 | 0.009 |
| Sex | F(1,62)=0.368 | 0.546 | 0.006 |
| Genotype × Sex | F(2,62)=0.706 | 0.497 | 0.022 |

Supplemental Table S15. Two-way ANOVAs of average of context fear expression during minutes 2-6. Refer to Supplemental Figures S2, S3.

| **Phase 1 Cued** | | | |
|---|---|---|---|
| | ***F statistic*** | ***p*** | ***partial $\eta^2$*** |
| Genotype | F(1,88)=0.006 | 0.938 | 0.000 |
| Sex | F(1,88)=1.056 | 0.307 | 0.012 |
| Genotype × Sex | F(1,88)=0.196 | 0.659 | 0.002 |

| **Phase 1 Context** | | | |
|---|---|---|---|
| | ***F statistic*** | ***p*** | ***partial $\eta^2$*** |
| **Genotype** | **F(1,77)=5.646** | **0.020** | **0.068** |
| Sex | F(1,77)=1.238 | 0.269 | 0.016 |
| Genotype × Sex | F(1,77)=0.646 | 0.424 | 0.008 |

| **Phase 2 Cued** | | | |
|---|---|---|---|
| | ***F statistic*** | ***p*** | ***partial $\eta^2$*** |
| Genotype | F(1,88)=0.006 | 0.938 | 0.000 |
| Sex | F(1,88)=1.056 | 0.307 | 0.012 |
| Genotype × Sex | F(1,88)=0.196 | 0.659 | 0.002 |

| **Phase 2 Context** | | | |
|---|---|---|---|
| | ***F statistic*** | ***p*** | ***partial $\eta^2$*** |
| Genotype | F(1,88)=0.006 | 0.938 | 0.000 |
| Sex | F(1,88)=1.056 | 0.307 | 0.012 |
| Genotype × Sex | F(1,88)=0.196 | 0.659 | 0.002 |

Supplemental Table S16. Two-way ANOVAs of fecal boli during swim stress. Refer to Supplemental Figure S4.

| **Phase 1 Cued** | | | |
|---|---|---|---|
| | ***F statistic*** | ***p*** | ***partial $\eta^2$*** |
| Genotype | F(1,45)=0.013 | 0.910 | 0.000 |
| Sex | F(1,45)=0.015 | 0.902 | 0.000 |
| Genotype × Sex | F(1,45)=0.604 | 0.441 | 0.013 |

| **Phase 1 Context** | | | |
|---|---|---|---|
| | ***F statistic*** | ***p*** | ***partial $\eta^2$*** |
| Genotype | F(1,36)=0.032 | 0.858 | 0.001 |
| Sex | F(1,36)=2.235 | 0.144 | 0.058 |
| Genotype × Sex | F(1,36)=0.122 | 0.729 | 0.003 |

| **Phase 2 Cued** | | | |
|---|---|---|---|
| | ***F statistic*** | ***p*** | ***partial $\eta^2$*** |
| Genotype | F(1,43)=1.661 | 0.204 | 0.037 |
| **Sex** | **F(1,43)=6.354** | **0.016** | **0.129** |
| Genotype × Sex | F(1,43)=0.042 | 0.839 | 0.001 |

| **Phase 2 Context** | | | |
|---|---|---|---|
| | ***F statistic*** | ***p*** | ***partial $\eta^2$*** |
| Genotype | F(1,34)=0.297 | 0.589 | 0.009 |
| Sex | F(1,34)=1.126 | 0.296 | 0.032 |
| Genotype × Sex | F(1,34)=1.126 | 0.296 | 0.032 |

Supplemental Table 17. Specific numbers of mice graphed in each Figure.

| **Figure 2: Phase 1 Cued** | **Females** | | **Males** | |
|---|---|---|---|---|
| | **Wildtype** | **Heterozygous** | **Wildtype** | **Heterozygous** |
| Training | 20 | 19 | 25 | 33 |
| Cued Expression Testing & Extinction Training | 20 | 19 | 25 | 33 |
| Extinction Retention Testing | 20 | 19 | 25 | 32 |
| Context Fear Expression | 20 | 19 | 23 | 30 |
| Cued Fear Renewal | 20 | 19 | 22 | 31 |

| **Figure 3: Phase 1 Context** | **Females** | | **Males** | |
|---|---|---|---|---|
| | **Wildtype** | **Heterozygous** | **Wildtype** | **Heterozygous** |
| Training | 20 | 18 | 20 | 23 |
| Context Fear Expression | 20 | 18 | 20 | 23 |

| **Figures 4 & 5: Phase 2 Cued** | **Females** | | **Males** | |
|---|---|---|---|---|
| Training | **Wildtype** | **Heterozygous** | **Wildtype** | **Heterozygous** |
| No Swim | 11 | 11 | 9 | 11 |
| Swim | 12 | 11 | 9 | 11 |
| Cued Expression Testing & Extinction Training | | | | |
| No Swim | 11 | 11 | 7 | 10 |
| Swim | 12 | 11 | 7 | 10 |
| Extinction Retention Testing | | | | |
| No Swim | 10 | 9 | 9 | 11 |
| Swim | 10 | 10 | 10 | 12 |
| Context Fear Expression | | | | |
| No Swim | 11 | 11 | 9 | 11 |
| Swim | 12 | 11 | 10 | 12 |
| Cued Fear Renewal | | | | |
| No Swim | 10 | 11 | 7 | 10 |
| Swim | 11 | 11 | 10 | 12 |

| **Figures 6 & 7: Phase 2 Context** | **Females** | | **Males** | |
|---|---|---|---|---|
| Training | **Wildtype** | **Heterozygous** | **Wildtype** | **Heterozygous** |
| No Swim | 9 | 10 | 9 | 9 |
| Swim | 10 | 9 | 9 | 10 |
| Context Fear Expression | | | | |
| No Swim | 9 | 8 | 9 | 9 |
| Swim | 8 | 9 | 9 | 10 |

| **Figure 8: Corticosterone** | **Females** | | **Males** | |
|---|---|---|---|---|
| Phase 1 Cued | **Wildtype** | **Heterozygous** | **Wildtype** | **Heterozygous** |
| No Swim | 10 | 8 | 12 | 15 |
| Swim | 10 | 10 | 15 | 16 |
| Phase 1 Context | | | | |
| No Swim | 10 | 8 | 10 | 11 |
| Swim | 10 | 10 | 10 | 11 |

| **Figure 9: Corticosterone** | **Females** | | **Males** | |
|---|---|---|---|---|
| Phase 1 Cued | **Wildtype** | **Heterozygous** | **Wildtype** | **Heterozygous** |
| No Swim | 11 | 11 | 10 | 11 |
| Swim | 12 | 11 | 12 | 12 |
| Phase 1 Context | | | | |
| No Swim | 9 | 10 | 9 | 9 |
| Swim | 10 | 9 | 9 | 10 |

| **Figures 10 & 11: Swim behaviors** | **Females** | | **Males** | |
|---|---|---|---|---|
| | **Wildtype** | **Heterozygous** | **Wildtype** | **Heterozygous** |
| Phase 1 Cued | 10 | 10 | 9 | 14 |
| Phase 1 Context | 10 | 10 | 9 | 11 |
| Phase 2 Cued | 12 | 11 | 12 | 11 |
| Phase 2 Context | 10 | 8 | 10 | 10 |

Supplemental Figure S1.

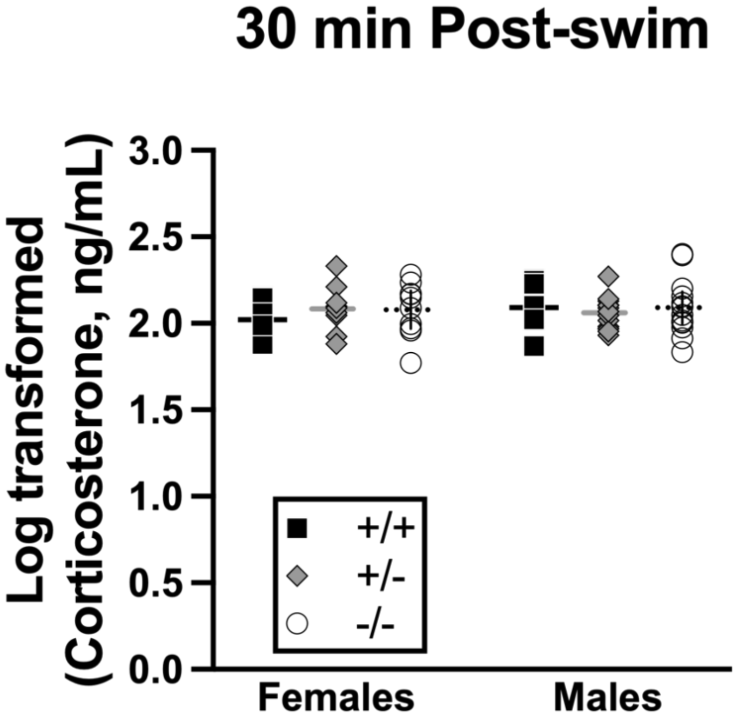


Supplemental Figure S1. ***Log-transformed corticosterone levels 30 min after an acute swim stress***.
Wildtypes are represented by black squares, heterozygotes by grey diamonds, and knockouts by clear circles. All mice underwent a six min swim stress, and blood was collected 30 min later to quantify serum corticosterone levels. Data are log-transformed cort levels [1–4]. Female wildtypes, n=10; female heterozygotes, n=9; female knockouts, n=10; male wildtypes, n=12; male heterozygotes, n=13, male knockouts, n=14. Data are graphed as mean (horizontal line) with 95% confidence interval (vertical line).

Supplemental Figure S2.

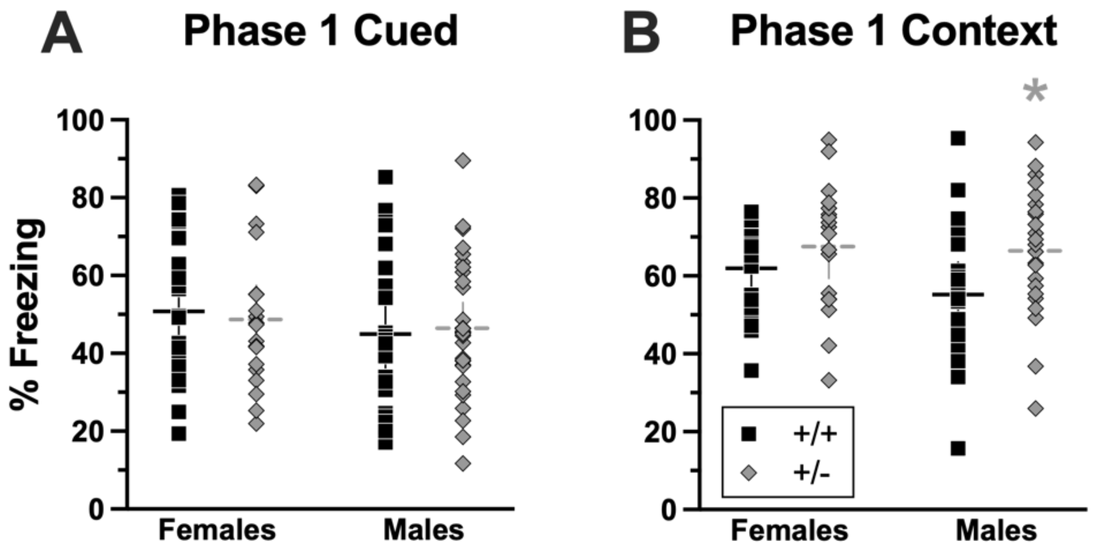


Supplemental Figure S2. ***Average context fear expression during minutes 2 through 6 of testing for Phase 1 mice***.
Wildtypes are represented by black squares, and heterozygotes are represented by grey diamonds. Context fear testing occurred on Day 5 for mice that underwent cued fear conditioning (A), whereas context fear testing occurred on Day 2 for mice that underwent context fear conditioning (B). Data are average percent time spent freezing during minutes two through six of the 10 min context fear testing period [2,5]. Phase 1 Cued: Female wildtypes, n=20; female heterozygotes, n=19; male wildtypes, n=23; male heterozygotes, n=30. Phase 1 Context: Female wildtypes, n=20; female heterozygotes, n=18; male wildtypes, n=20; male heterozygotes, n=23. Data are graphed as mean (horizontal line) with 95% confidence interval (vertical line). *p=0.0455 indicates difference between heterozygous and wildtype within same Phase 1 Context sex.

Supplemental Figure S3.

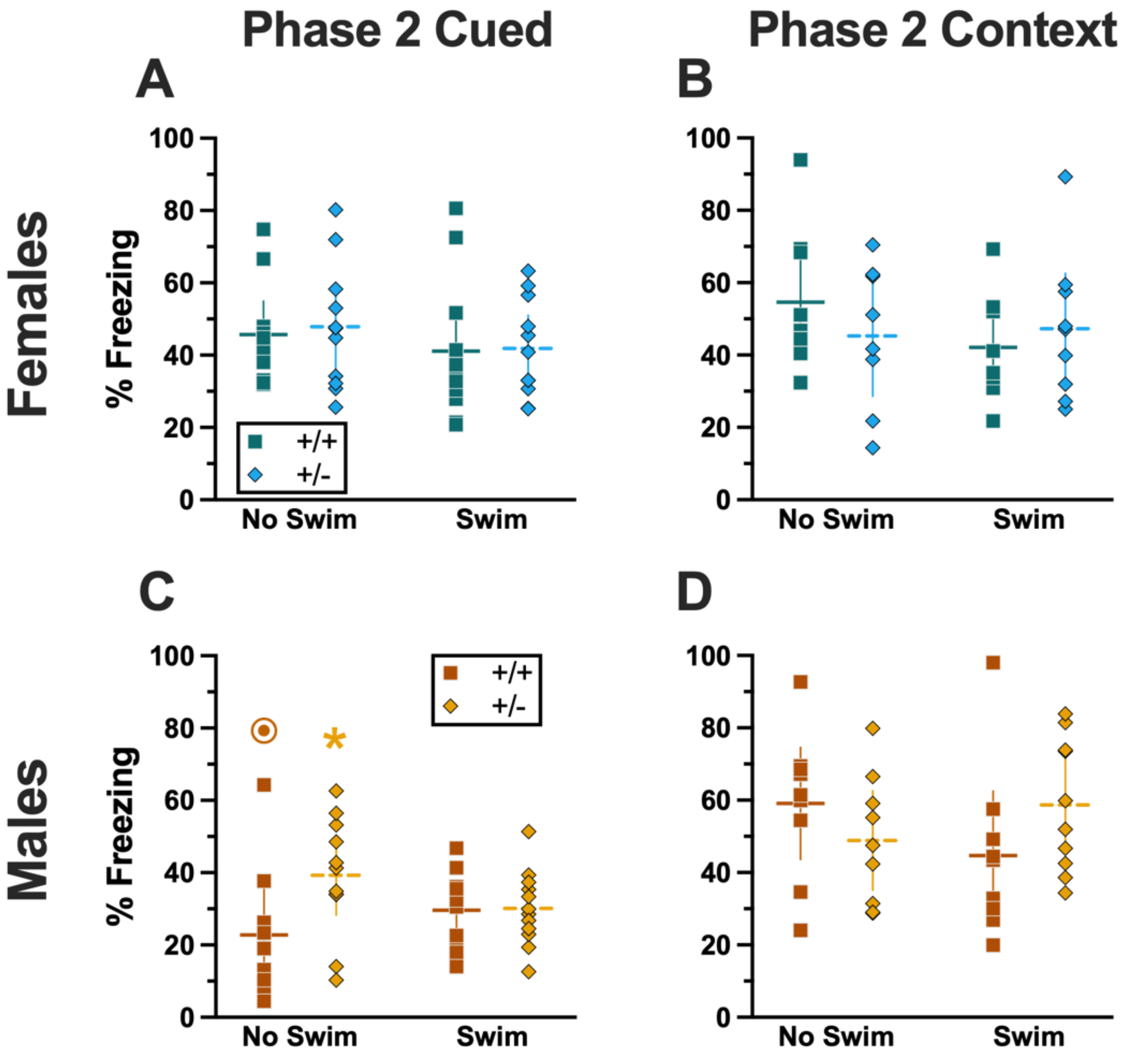


Supplemental Figure S3. ***Average context fear expression during minutes 2 through 6 of testing for Phase 2 mice***.
Female (A-B) wildtypes are represented by teal squares, and female heterozygotes are represented by blue diamonds. Male (C-D) wildtypes are represented by orange squares, and male heterozygotes are represented by yellow diamonds. Context fear testing occurred on Day 5 for mice that underwent cued fear conditioning (A,C), whereas context fear testing occurred on Day 2 for mice that underwent context fear conditioning (B,D). Data are average percent time spent freezing during minutes two through six of the 10 min context fear testing period [2,5]. Phase 2 Cued No Swim: Female wildtypes, n=11; female heterozygotes, n=11; male wildtypes, n=9; male heterozygotes, n=11. Phase 2 Cued Swim: Female wildtypes, n=12; female heterozygotes, n=11; male wildtypes, n=10; male heterozygotes, n=12. Phase 2 Context No Swim: Female wildtypes, n=9; female heterozygotes, n=8; male wildtypes, n=9; male heterozygotes, n=9. Phase 2 Context Swim: Female wildtypes, n=8; female heterozygotes, n=9; male wildtypes, n=9; male heterozygotes, n=10. Data are graphed as mean (horizontal line) with 95% confidence interval (vertical line). *p=0.019 indicates difference between

heterozygous and wildtype within same sex and swim condition. ⊙p=0.001 indicates difference between sexes within same genotype and swim condition.

Supplemental Figure S4.

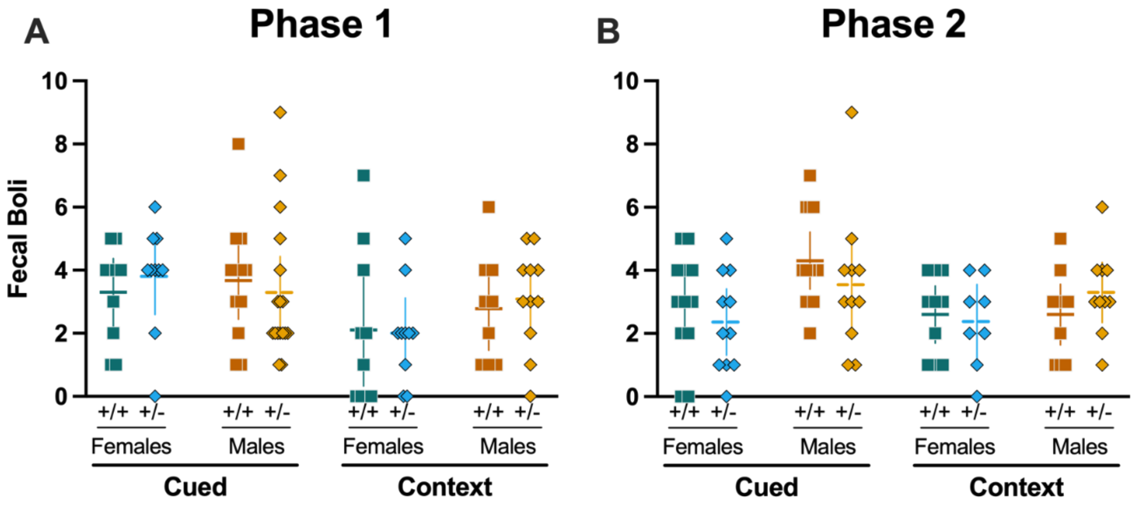


Supplemental Figure S4. ***Fecal boli from swim stressors across Phases***.
Female wildtypes are represented by teal squares, and female heterozygotes are represented by blue diamonds. Male wildtypes are represented by orange squares, and male heterozygotes are represented by yellow diamonds. Mice were exposed to six minutes of swim stress after (Phase 1) or before (Phase 2) cued or context fear conditioning. Data are fecal boli produced during swim stress. Phase 1 Cued: Female wildtypes, n=10; female heterozygotes, n=10; male wildtypes, n=12; male heterozygotes, n=17. Phase 1 Context: Female wildtypes, n=10; female heterozygotes, n=10; male wildtypes, n=9; male heterozygotes, n=11. Phase 2 Cued: Female wildtypes, n=12; female heterozygotes, n=11; male wildtypes, n=13; male heterozygotes, n=11. Phase 2 Context: Female wildtypes, n=10; female heterozygotes, n=8; male wildtypes, n=10; male heterozygotes, n=10. Data are graphed as mean (horizontal line) with 95% confidence interval (vertical line).

Excluded fear behavior data

| Phase 1 Cued | Reason |
|---|---|
| 0608 | All fear data excluded because cued fear expression never exceeds 25% |
| 0609 | Data are excluded for context fear testing & cued fear renewal because camera was laggy for 609, 610, and 616, thus freezing data are inaccurate |
| 0610 | Data are excluded for context fear testing & cued fear renewal because camera was laggy for 609, 610, and 616, thus freezing data are inaccurate |
| 0616 | Data are excluded for context fear testing & cued fear renewal because camera was laggy for 609, 610, and 616, thus freezing data are inaccurate |
| 0664 | First 5 tones of cued fear expression <25%, so all data excluded |
| 0681 | All fear data excluded because cued fear training never exceeds 25% |
| 0768 | First 5 tones of cued fear expression <25%, so all data excluded |
| 0770 | No freezing during context testing or cued fear renewal>25% |
| 1057 | No freezing during context testing >25%, context testing data excluded |
| 1182 | All fear data excluded because cued fear training never exceeds 25% |
| 1168 | Cued fear renewal data excluded because cued fear renewal never exceeds 25% |
| 0828 | First 5 tones of cued fear expression <25%, so all data excluded |
| 1255 | First 5 tones of cued fear expression <25%, so all data excluded |
| 1264 | First 5 tones of cued fear expression <25%, so all data excluded |
| 1284 | First 5 tones of cued fear expression <25%, so all data excluded |

| Phase 1 Context | Reason |
|---|---|
| 0713 | Shock did not work, was accidentally turned off; exclude all data because never trained |
| 0771 | Excluding data because mouse exhibited >75% freezing prior to first shock |
| 0775 | Excluding data because mouse exhibited >75% freezing prior to first shock |
| 0776 | No freezing during testing >25%, all data excluded |
| 0832 | No freezing during testing >25%, all data excluded |
| 1081 | Exceeded 75% freezing prior to first shock being administered; excluding all data |
| 1070 | No freezing during testing >25%, all data excluded |
| 1080 | No freezing during testing >25%, all data excluded |

| Phase 2 Cued | Reason |
|---|---|
| 0782 | All data excluded because first 5 tones cued fear expression on first testing day never exceed 25% |
| 0792 | Cued fear renewal data excluded because cued fear renewal never exceeds 25% |
| 0796 | Cued fear renewal data excluded because cued fear renewal never exceeds 25% |
| 0827 | Mice were not run under correct protocol for testing day 2, so excluding data from this testing day (were run under context testing/cued fear renewal protocol, though in context B) |
| 0869 | Context testing & cued fear renewal testing, door was open and mouse left the chamber at 532.67, data from 510 s onwards on this day excluded |
| 0870 | Cued fear renewal data excluded because cued fear renewal never exceeds 25% |
| 0873 | Mice were not run under correct protocol for testing day 2, so excluding data from this testing day (were run under context testing/cued fear renewal protocol, though in context B) |
| 0879 | Mice were not run under correct protocol for testing day 2, so excluding data from this testing day (were run under context testing/cued fear renewal protocol, though in context B) |
| 0881 | Videos did not record for cued expression testing & extinction training (7 May 2022) for these animals, so must exclude from analyses |
| 0885 | Mice were not run under correct protocol for testing day 2, so excluding data from this testing day (were run under context testing/cued fear renewal protocol, though in context B) |
| 0886 | Mice were not run under correct protocol for testing day 2, so excluding data from this testing day (were run under context testing/cued fear renewal protocol, though in context B) |
| 0887 | Mice were not run under correct protocol for testing day 2, so excluding data from this testing day (were run under context testing/cued fear renewal protocol, though in context B) |
| 0908 | Videos did not record for cued expression testing & extinction training (7 May 2022) for these animals, so must exclude from analyses |
| 0909 | Videos did not record for cued expression testing & extinction training (7 May 2022) for these animals, so must exclude from analyses |
| 0910 | Videos did not record for cued expression testing & extinction training (7 May 2022) for these animals, so must exclude from analyses |
| 0916 | Videos did not record for cued expression testing & extinction training (7 May 2022) for these animals, so must exclude from analyses |
| 0917 | Videos did not record for cued expression testing & extinction training (7 May 2022) for these animals, so must exclude from analyses; Cued fear renewal data excluded because cued fear renewal never exceeds 25% |
| 0918 | Videos did not record for cued expression testing & extinction training (7 May 2022) for these animals, so must exclude from analyses |
| 0919 | Videos did not record for cued expression testing & extinction training (7 May 2022) for these animals, so must exclude from analyses |

| | |
|---|---|
| 0923 | All data excluded because cued fear expression on first testing day never exceeds 25% |
| 0924 | Training video did not save/record; All data excluded because cued fear expression on first testing day never exceeds 25% |
| 0925 | Training video did not save/record |
| 0929 | Training video did not save/record; All data excluded because cued fear expression on first testing day never exceeds 25% |
| 0944 | Training video did not save/record |
| 0945 | All data excluded because cued fear expression on first testing day never exceeds 25% |
| 0946 | All data excluded because cued fear expression on first testing day never exceeds 25% |
| 0953 | Exclude all fear data. These animals were swam in both cohorts 19 and 20. Only Swim scoring of cohort 19 will be used. |
| 0967 | Exclude all fear data. These animals were swam in both cohorts 19 and 20. Only Swim scoring of cohort 19 will be used. |
| 0981 | All data excluded because cued fear expression to first 5 tones on first testing day never exceeds 25% |
| 1192 | All data excluded because first 5 tones cued fear expression on first testing day never exceed 25% |
| 1210 | All fear data excluded because cued fear training never exceeds 25% |
| 1218 | Cued fear renewal data excluded because cued fear renewal never exceeds 25% |
| 1212 | All data excluded because cued fear expression to first 5 tones on first testing day never exceeds 25% |
| 1223 | All fear data excluded because cued fear training never exceeds 25% |

| **Phase 2 Context** | **Reason** |
|---|---|
| 1083 | All data excluded because context fear expression never exceeds 25% |
| 1092 | Testing data didn't record because hard drive was not cleared on schedule |
| 1098 | Testing data didn't record because hard drive was not cleared on schedule |
| 1079 | Testing data didn't record because hard drive was not cleared on schedule |
| 1106 | Testing data didn't record because hard drive was not cleared on schedule |
| 1141 | Exceeded 75% freezing prior to first shock being administered; excluding all data |

Excluded swim data

| Male・Phase 1 Context・Wildtype<br>Climbing Behavior・(All Swim Data Excluded for 0836) | | | |
|---|---|---|---|
| **Animal ID** | **Data** | **Outlier Determination** | |
| 0700 | 44.1 | | |
| 0723 | 0.0 | 22.0 | *Avg* |
| 0726 | 16.6 | 16.7909946 | *SD* |
| 0783 | 28.5 | 105.92164 | *Avg + 5 SD* |
| 0819 | 4.6 | | |
| 0820 | 0.9 | | |
| 0836 | 121.5 | >5 SD beyond mean | |
| 0857 | 36.4 | | |
| 1120 | 34.0 | | |
| 1146 | 32.6 | | |

| Male・Phase 2 Cued・Wildtype<br>Swimming Behavior・(All Swim Data Excluded for 0944) | | | |
|---|---|---|---|
| **Animal ID** | **Data** | **Outlier Determination** | |
| 0881 | 119.9 | | |
| 0909 | 177.1 | | |
| 0919 | 138.1 | | |
| 0944 | 356.9 | >8 SD beyond mean | |
| 0946 | 172.1 | | |
| 0952 | 118.3 | | |
| 0967 | 159.4 | | |
| 1009 | 151.4 | | |
| 1034 | 132.4 | 135.1 | *Avg* |
| 1036 | 104.2 | 25.7674959 | *SD* |
| 1038 | 130.0 | 341.269134 | *Avg + 8 SD* |
| 1045 | Recording failed | | |
| 1212 | 124.8 | | |
| 1223 | 93.9 | | |

| Male・Phase 2 Cued・Heterozygous<br>Climbing Behavior・(All Swim Data Excluded for 0869) | | |
|---|---|---|
| **Animal ID** | **Data** | **Outlier Determination** |
| 0869 | 153.3 | >5 SD beyond mean |
| 0908 | 4.0 | |
| 0910 | 11.1 | |
| 0899 | 22.8 | |
| 0904 | 63.2 | |

| | | | |
|---|---|---|---|
| 0925 | 34.1 | | |
| 0935 | 32.5 | | |
| 0994 | 11.5 | | |
| 1007 | 22.3 | 33.6 | *Avg* |
| 1016 | 31.6 | 23.2230254 | *SD* |
| 1086 | 67.2 | 149.706036 | *Avg + 5 SD* |
| 1125 | 69.2 | | |

Female・Phase 2 Context・Heterozygous
Climbing Behavior・(All Swim Data Excluded for 0955)

| Animal ID | Data | Outlier Determination | |
|---|---|---|---|
| 0921 | 25.9 | | |
| 0922 | 4.4 | 14.4 | *Avg* |
| 0937 | 0.5 | 15.7784437 | *SD* |
| 0943 | 44.5 | 140.65255 | *Avg + 8 SD* |
| 0949 | 1.4 | | |
| 0955 | 144.0 | >8 SD beyond mean | |
| 1023 | 25.4 | | |
| 1092 | 7.4 | | |
| 1158 | 5.9 | | |
